# E-Protein Protonation Titration-induced Single Particle Chemical Force Spectroscopy for Microscopic Understanding and pI Estimation of Infectious DENV

**DOI:** 10.1101/2023.03.30.534862

**Authors:** Manorama Ghosal, Tatini Rakhshit, Shreya Bhattacharya, Sankar Bhattacharyya, Priyadarshi Satpati, Dulal Senapati

## Abstract

The ionization state of amino acids on the outer surface of a virus regulates its physicochemical properties toward the sorbent surface. Serologically different strain of dengue virus (DENV) shows different extents of infectivity depending upon their interactions with a receptor on the host cell. To understand the structural dependence of E-protein protonation over its sequence dependence, we have followed E-protein titration kinetics both experimentally and theoretically for two differentially infected dengue serotypes, namely DENV-2 and DENV-4. We have performed an E-protein protonation titration-induced single particle chemical force spectroscopy using an atomic force microscope (AFM) to measure the surface chemistry of DENV in physiological aqueous solutions not only to understand the charge distribution dynamics on virus surface but also to estimate the isoelectric point (pI) accurately for infectious dengue viruses. Cryo-EM structure-based theoretical pI calculations of DENV-2 surface protein were shown to be consistent with the evaluated pI value from force spectroscopy measurements. This is a comprehensive study to understand how the cumulative charge distribution on the outer surface of a specific serotype of DENV regulates a prominent role of infectivity over minute changes at the genetic level.

## INTRODUCTION

Dengue virus (DENV) belongs to the flavivirus family which is most commonly known for dengue hemorrhagic fever (DHF) and dengue shock syndrome (DSS) worldwide. About 3.6 billion people are at risk and 390 million infections happen annually where 96 million are symptomatic cases with considerable lethality^1^. DENV infections prevail in developing and underdeveloped countries where the surveillance network for infectious diseases is not strong enough and thus the possibility of gross under-reporting of dengue is high. It is also important to note that since 1779 DENV infections have been dominating many areas of Asia and the Pacific regions. The major epidemics occurred in the Philippines and Greece in 1920 and the South Pacific in 1930^2,3^. Though only one version of the dengue vaccine, Dengvaxia, is commercially available in the market, at best 20 countries have licensed it due to its low potency toward different serotypes of DENV^4^. Many antiviral agents have been designed by targeting structural and non-structural proteins of DENV, but maximum findings are stopped in the pre-clinical research pipeline due to low efficacy and pharmacokinetic uncertainty. Researchers are mainly aiming for the E protein as the target area of DENV for effective drug design to avoid intercellular toxicity and membrane permeability^5^. The E protein is composed of three conserved functional domains: (1) Flavivirus envelope glycoprotein E, stem/anchor domain, (2) Flavivirus glycoprotein, central and dimerization domains, and (3) Flavivirus glycoprotein, immunoglobulin-like domain. Out of three domains (E-I, E-II, and E-III) in the E protein, the E-III domain is characterized by an immune globulin-like structure containing the most distal projecting loops from the virion surface that facilitates the binding of viral particles with the cell surface^6,7^. After viral particles enter the cell through the clathrin-coated endocytic pathway, there must be a conformational change of viral E protein from dimer to trimer, depending upon the lower pH environment of the endosome^8,9^. This trimeric E protein then opens up its E-II domain which contains a fusion loop for disruption of the endosomal membrane and releases the viral nucleocapsid into the host cytosol for further expression of viral RNA^10^. Thus, DENV E protein-based (mainly E-III) agents are aimed by so many researchers as antiviral drugs. However, due to its low potency, this detection could not find its interference up to clinical uses. As the structural integrity of the E protein is induced by the pH of the surroundings, the total surface charge of DENV will be changed depending on the protonation state of the E proteins^11^. The attraction and repulsion between the virus and any charged surface are mainly governed by short and long-range forces, this phenomenon is modulated by the pH and ionic strength of the solution^12^. Other than electrostatic forces, van der Waals forces, hydrophobic interactions, and H-bonds are also crucial for specific binding and nucleocapsid release of DENV. Many studies have shown evidence that E protein promoted viral entry^13^, but how the viral envelope changed the total surface charge of the DENV surface depending upon the H^+^ concentration of the surrounding is still missing. It is also poorly understood how the virus surface charges are governed by the ionizable amino acids on the outer membrane part of the E protein and other factors like virus capsids and negatively charged nucleic acid within the core of the capsid^14^. The surface polarity of DENV will help us to describe its physicochemical properties linked with its adsorption and maturation. The immobilization of DENV on the host cell surface is largely through ionic interactions between the virion particles and oppositely charged components of the host cell extracellular matrix. Such interactions increase the local concentration of the virus on host cells increasing the chances of an interaction with the cognate entry receptor. Subsequently, the receptor-bound virus particle is internalized into an endosome, which is followed by acidification of the endosome lumen by proton pumps on the endosome membrane^15^. Upon reduction of pH, the dimeric form of E protein converts to a trimeric form, a transition necessary for the fusion of the viral and endosomal membrane, which is followed by the release of the viral genomic RNA into the host cell. Though the attachment of DENV on target cell surface receptors is governed by the charged amino acids of E-III^16^, it is also known that the endosomal fusion of DENV is mainly commanded by the three hydrophobic amino acids (W101, L107, and F108) within the fusion loop of trimeric E protein^17^. So, the tropism of DENV is highly influenced by charge densities of the specific domains of the E protein which folded up differentially at variable pH. Among 4 different serotypes of DENV, the infectiousness is dissimilar as little genetic variance tends to a large variation within the structural orientation of the active pockets of the proteins. This kind of structural differentiation of E protein can be measured by looking into the cumulative surface charge of DENV in different protonation states of E glycoprotein at a single particle level. The surface charge of DENV can be used to deduce the adhesion forces between the virus and its charged target substrate to elicit its attachment behavior^16^. The surface charge of any particle plays a characteristic conversion with respect to the protonation state of its charged residues. At a certain pH, where the net collective surface charge is zero, known as the isoelectric point (pI) of the particle. Likely, when DENV is in any certain pH solution then its charged residues interact with the solvent molecules to form an electrical bilayer which we can deduce by measuring the zeta potential, isoelectric focusing, or other bulk methods. But, these methods are complicated concerning sample purification procedures and disruption of the intact virus^18,19, 20^. Thus, we have preferred a novel surface characterization method known as chemical force microscopy (CFM), using an atomic force microscope (AFM) to measure the surface charge distribution of DENV in physiological aqueous solutions^21,22^. A total of 90 head-tail dimers of E-proteins cover the outermost part of the DENV to form a protein shell of the virus. Thus, in a single particle study at different pH, the accurate adhesion forces between the virus surface and the charged AFM probe are mostly arising from different charge distributions of the E protein^8^. In principle, a theoretical titration curve can be obtained by determining the pH-dependent protonation states of the E protein extracellular domain. Replication of the theoretical E-Protein Protonation Titration (EPPT) can be complemented by a single-particle chemical force spectroscopy (SPCFS) for microscopic understanding and accurate estimation of charge states for different infectious DENV serotypes. CFS with chemically modified probes of a cantilever can map and quantify chemical heterogeneities on the single particle surface ^23,24^. The surface charge of a single *Saccharomyces cerevisiae* and the pI value of various proteins have been evaluated by CFM^25^. There are several other noted microbiological studies that are also being reported in the literature by using CFM as a path-breaking *in situ* microscopic technique. Viljoen et al. has shown the importance of nanodomain hydrophobicity for the pathogenesis of *Mycobacterium abscessus*^26^. Guinlet et al. have reported how an adequate force controls the colonization and infection of gram-positive viruses^27^. Very recently, Koehler et al. have used single particle CFM to find out the interaction strategy of reovirus with β1 integrins for cell internalization in a clathrin-mediated process^28^. Many other ligand-receptor interactions, such as antibody-antigen interaction and protein-protein interaction are elucidated by the force spectroscopy technique^22,29,30^. The advantage of CFM over other methods lies in its ability to perform in physiological conditions where the structural integrity of the virus is not hampered and very little sample volume is enough for consecutive experiments^31, 32^. It is the best of our knowledge that no single-vesicle level isoelectric point determination was carried out before on any human flavivirus samples.

Here we have taken two DENV serotypes, DENV-2 and DENV-4, to evaluate their net surface charge at different pHs of the medium. DENV-2 is considered to be the most prevalent serotype, whereas DENV-4 has comparatively lower extremities^33,34^. It has been seen that the phylogenetic analysis to determine the infectivity of DENV strains in Asiatic and Pacefic regions always showed a comparable less severity for DENV-4, thus we have tried to get an idea about their surface charge distribution against a very infectious strain DENV-2 (Serotype influences on dengue severity: a cross-sectional study on 485 confirmed dengue cases in Vitória, Brazil). We have measured the adhesion force on the DENV surface with a functionalized (with either a positive or a negative charge terminated organic molecule) AFM probe^32^ and evaluated the pI of DENV i.e., the value of pH where the net surface charge of the specified serotypes are zero. We have compared the pI values obtained from detailed CFM studies with the zeta potential measurements and computationally verified the protonation state of the envelope protein as a function of pH for both DENV-2 and DENV-4. We have determined the surface pI of mammalian infectious virus by CFM for the first time and also computationally showed the distribution of charges on the outer surface of the E protein and around the fusion loop of the E trimer. This method will help us further to understand how the modulation of surface charge triggers the infection and maturation pathways of DENV and other enveloped viruses at a single particle level.

## RESULT AND DISCUSSION

### Adsorption of the virus on a mica surface and probe modification

A detailed experimental section which includes the preparation of different buffers (**Table S1**), Inactivation of virus samples, Functionalization procedure of AFM probe by different charged chemical groups (**Figures S1** and **S2**), AFM imaging and data analysis (**Figure S3** and **S4**) by Chemical Force Measurement (CFM), Zeta potential measurement of inactivated DENV of all serotypes, and the associated computational methodology for the estimation of pI (**Table S2** and **Figure S5-S8**) has been elaborated in the Supplementary Information (**SI**) section. A V1-type muscovite spherical mica sheet was first coated with 3-APTES ((3-aminopropyl triethoxysilane). When free-OH groups on the mica surface react with 3-APTES, it gives a smooth and hydrophobic surface compared to the bare mica surface. Viruses or any other microorganisms can easily be then adsorbed on such a solid surface hydrophobically from the gain in free energy for the systems as their adsorption minimizes the interfacial area between the apolar surface of the virus and buffer^12^. The deposited virus concentration on 3-APTES-treated mica was optimized to get a well-dispersed film for force spectroscopy. The obtained film was stable enough to avoid any sort of sample trip-off during CFM experiments as shown in the topographic image in **Figure 1a** and its corresponding 3D image in **Figure 1c**. The height profile of a single virus (cyan line) was measured from **Figure 1a** and then plotted in **Figure 1b**. The measured average height of the viruses is 18 ± 2 nm and the diameter of the virus is the distance between two dotted lines within **Figure 1b**. The diameter of the twenty-five different DENV2 have been measured in liquid mode and the obtained average diameter is ~ 72 nm. All CFM measurements were carried out on the virus film to increase the reproducibility of measurements.

**Figure 1:**
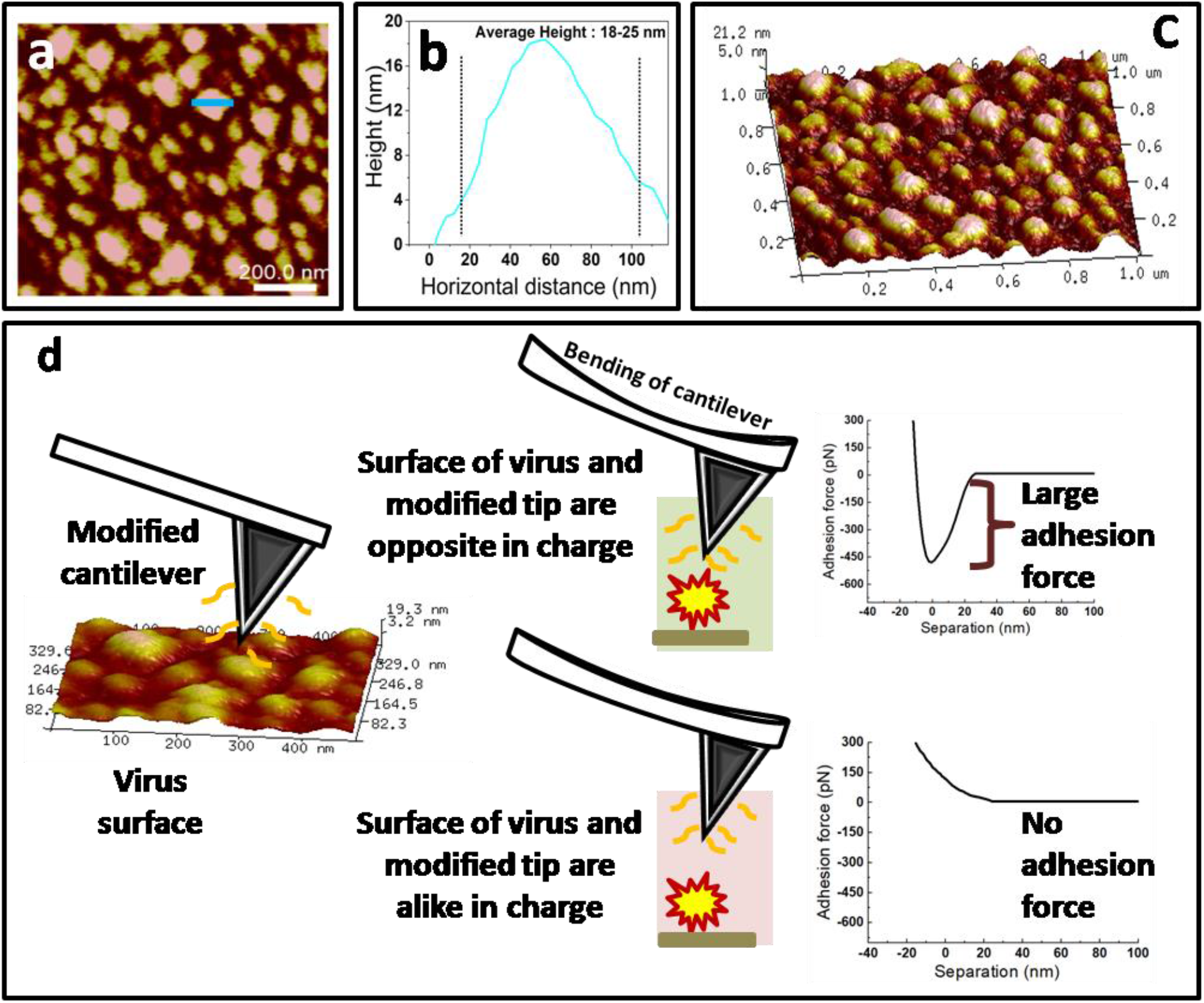
(a) Topographic image of DENV film under buffer, (b) height profile of a single virus, (c) corresponding 3D image of the DENV film as shown in **Figure 1(a)**, and (d) schematic representation of the type of interactions depending on the similarity and dissimilarity of charges between the surface and approaching tips during CFM measurements.

We used citrate buffer at various pH with positive or negative charge-modified tips. The tip modification was done in 2 steps. In the 1^st^ step, the tip was altered with 3-MPTMS to get a hanging mercapto (-S-H) group on the surface and in the 2^nd^ step, this mercapto group makes a - S-S-bond with positive 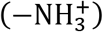 or negative (COO^-^) charge terminated molecules as shown in **Figure S1**^35^. The surface charge of the virus is highly dependent on the pH of the surroundings. The negative charge is enhanced as the pH is moved from a lower to a higher value and thus different amounts of adhesion force can be generated between the surface of a specific DENV serotype and the charged tip. **Figure 1d** schematically demonstrates how the adhesion force changes depending on the similarity and dissimilarity of charges between the DENV surface and approaching tips during CFM measurements.

### The adhesion of virus surface with differently modified probe using CFM

The DENV film interacts differently with a modified tip in presence of a buffer with various pHs (from 2.2 to 6.0). When the DENV surface and probes are opposite in charge, approaching the tip on the surface of the DENV maximizes the adhesion due to the electrostatic attraction between them. On the contrary, when both are similar in charge, an almost zero adhesion was observed due to electrostatic repulsion between them. Virus surface proteins are not fully protonated or deprotonated in any pH state between 2.2 to 6.0 and thus the adhesion forces were gained in a range that we plotted as a histogram. The negative charge-terminated probe was first approached on the DENV-2 film in the presence of 20 mM citrate buffer (CB) with a pH of 2.2, where we observed the maximum adhesion force as shown in **Figure 2a**. Once we increase the pH further, the adhesion force reduces gradually due to the gradual enhancement of DENV-2 surface negativity. The measured adhesion force was almost negligible when the pH reached 6.0 due to a strong repulsion between the alike-charged DENV-2 and the charge-terminated probe. Similar force measurements were carried out with a positive charge-terminated probe at different pHs in the range of 2.2-6.0 and we observed an opposite effect as observed for the negative charge-terminated probe. The corresponding adhesion force statistics are shown in **Figure 2c**. The positive probe adhered most strongly with the DENV-2 surface when the pH is 6 and the adhesion force reduces gradually from 6.0 to 2.2. The corresponding force-distance (F-D) curves are shown in **Figures 2b** and **2d** for negative- and positive-charged probes respectively. The same set of experiments was performed for DENV-4 too, where the tendency of adhesion histograms show a close correlation with DENV-2. For DENV-4, **Figures 3a** and **3c** show adhesion histograms for negative- and positive-charged probes, respectively, and the corresponding force-distance curves are shown in **Figures 3b** and **3d**, respectively. Worth to mention here that the DENV film was comparatively unstable below pH 3 for both serotypes.

**Figure 2:**
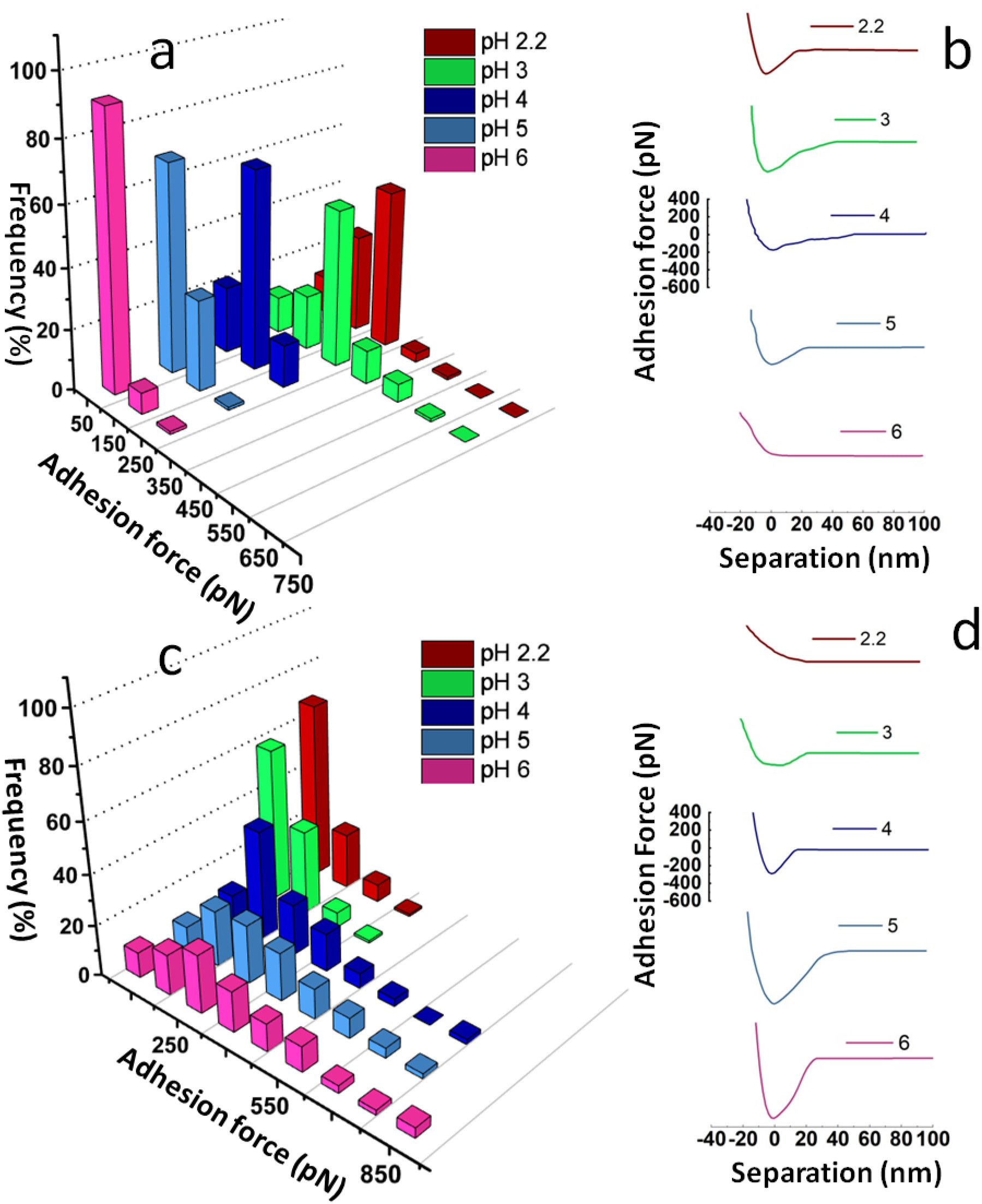
Adhesion histograms and representative force-distance curves: Adhesion force (pN) vs. frequency (%) for DENV-2 with (a) negative- and (c) positive-charge terminated probes with varying pH at 2.2, 3, 4, 5, and 6. Corresponding retract curves for (b) negative- and (d) positive-charge terminated probes with the same varying pH at 2.2, 3, 4, 5, and 6.

**Figure 3.**
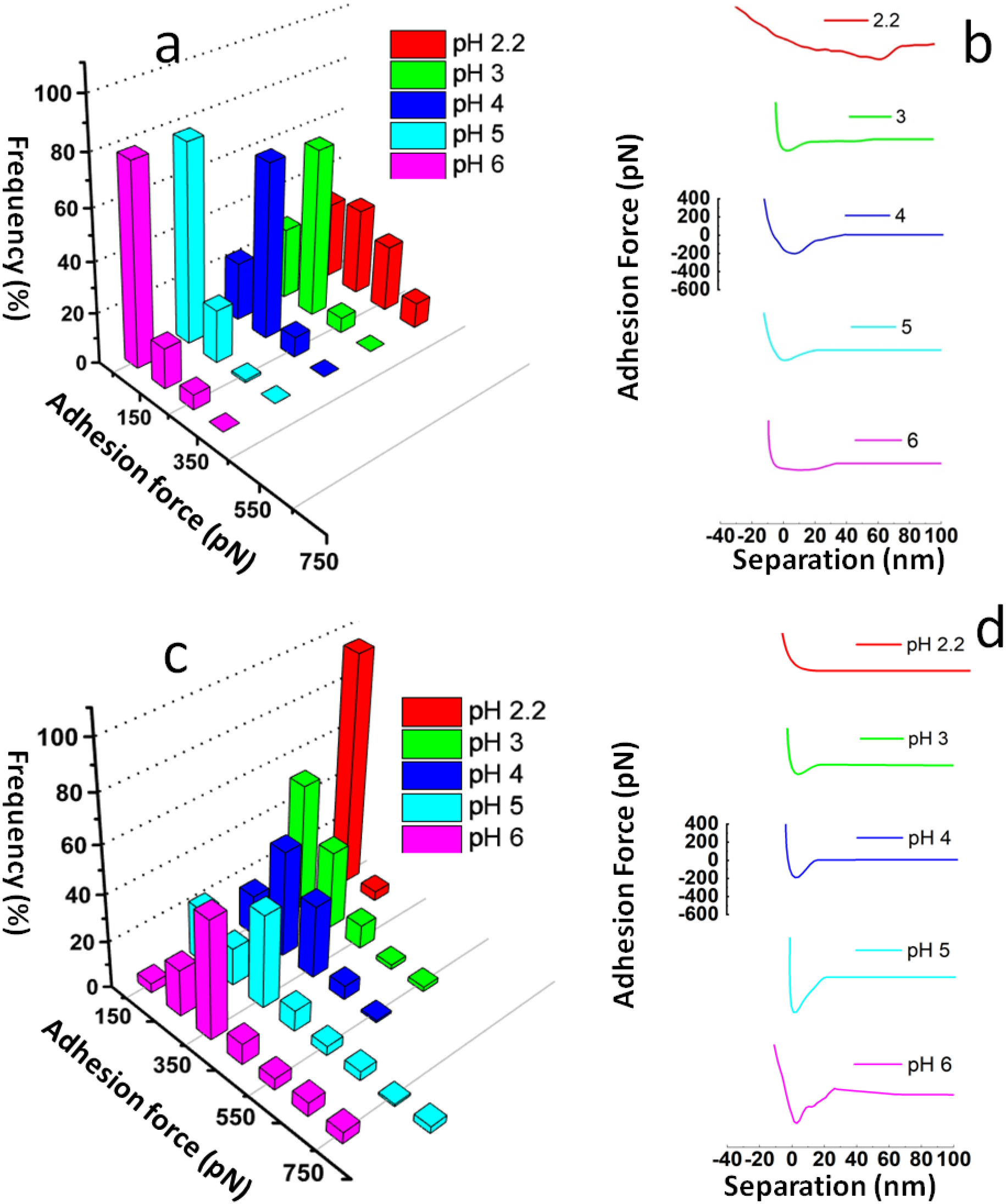
Adhesion histograms and representative force-distance curves: Adhesion force (pN) vs. frequency (%) for DENV-4 with (a) negative- and (c) positive-charge terminated probes with varying pH at 2.2, 3, 4, 5, and 6. Corresponding retract curves for (b) negative- and (d) positive-charge terminated probes with the same varying pH at 2.2, 3, 4, 5, and 6.

### Accurate Estimation of pI for Infectious DENV

As shown in **Figures 2** and **3**, a range of forces (between 0-750 pN) is statistically distributed on the adhesion histograms at different pHs. For a particular pH, the mean value of the adhesion force has then been calculated by considering more than 500 F-D curves, and the standard deviation of pH-dependent adhesion force is calculated by considering 3 sets of similar experiments at that particular pH. Obtained adhesion force against pH is then plotted for both negative- (**Figures 4b** and **4e**) and positive-charged tip (**Figures 4c** and **4f**). Finally, data points of the adhesion force vs. pH plot for each different type of DENV serotype were further fitted with equation (1) to get the sigmoidal plot.^36,32^

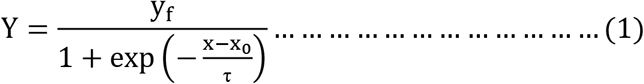

**Figure 4:**
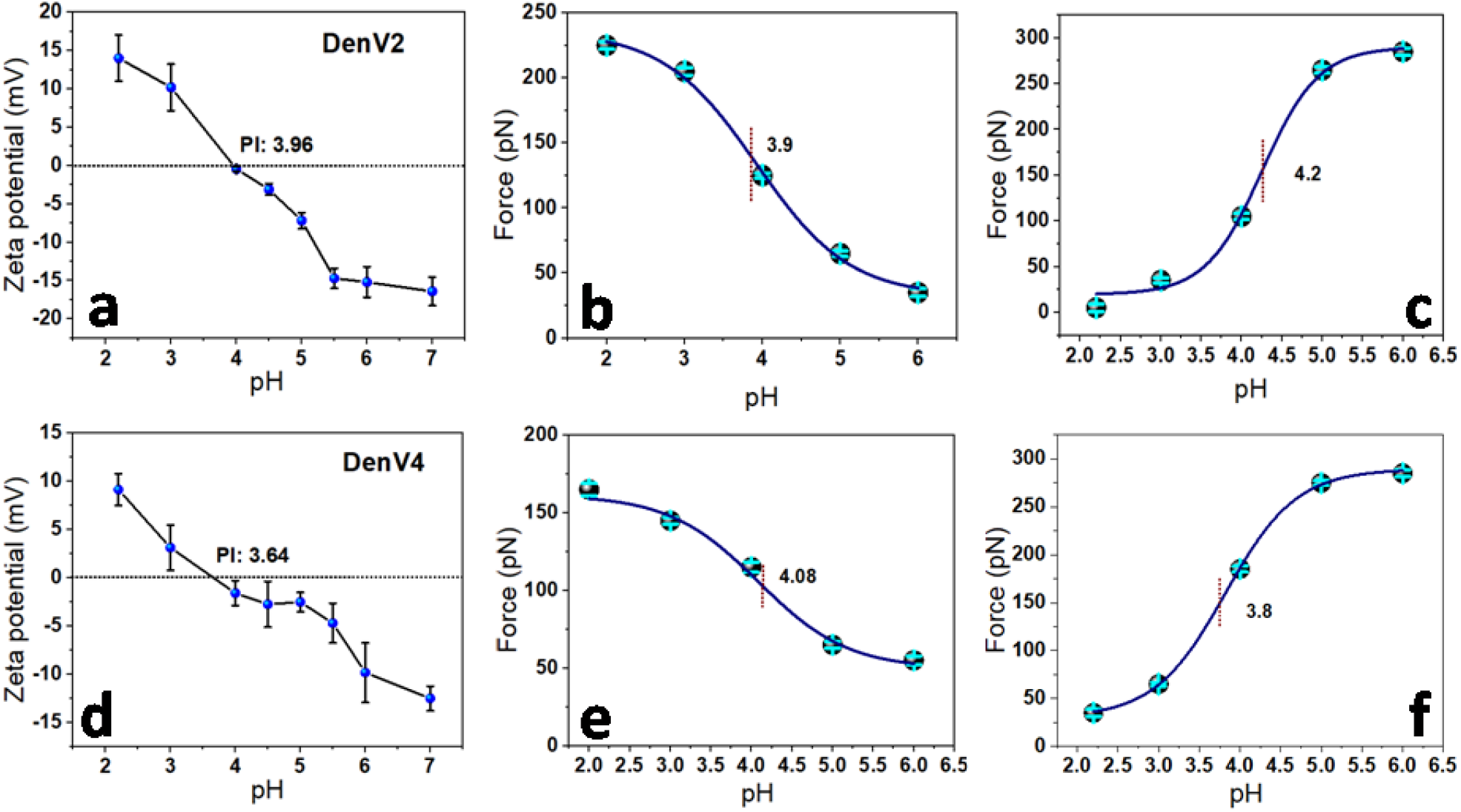
Estimation of the isoelectric point (pI): Variation of zeta potential with pH for (a) DENV-2, and (d) DENV-4; Variation of average adhesion force with pH for DENV-2, and DENV-4 by using a (b, e) negative-, and a (c, f) positive-charged probe respectively. The obtained data points from chemical force microscopy (CFM) at different pH are fitted with a modified Boltzmann sigmoidal formula. The standard deviations of force-pH curves are indicated with cyan color.

The pI of the DENV was measured from the inflection point of the obtained sigmoidal plots where Y and X are the adhesion force and corresponding pH respectively, yf is the maximum adhesion force, 1/τ is the rate constant for the change of the mean adhesion force as a function of pH and x0 is the pI which has been deduced from the plot. The obtained inflection points of the Force vs. pH curve from **Figures 4b and 4c** for DENV-2 are 3.9 and 4.2 with negative- and positive-charged probes respectively. By averaging these inflection points, the estimated pI for DENV-2 was recorded as 4.05. In the same way, inflection points obtained for DENV-4 are 4.08 and 3.8 from **Figures 4d and 4e,** giving the estimated pI for DENV-4 as 3.94.

To compare the pI value obtained from CFM with another commonly used method, we have adopted zeta potential as another technique. Zeta potentials of both the DENV serotypes were measured at various pHs as plotted in **Figure 4**. It has been observed that the zeta value shifts from positive to negative at pH 3.96 and 3.64 for DENV-2 (**Figure 4a**) and DENV-4 (**Figure 4d**), respectively. Apparently, these are the two pH values for DENV-2 and DENV-4, respectively, where the acquired surface charge is zero i.e., the pI for respective serotypes. Though the described CFM method monitors the pH-dependent resultant charge of the DENV surface at a single molecule level, zeta potential (or isoelectric focusing, IEF) is a bulk method where the surface charge is a double layer potential of suspended particle which is generally a little far from the actual surface. As a result, we can’t expect to get the same pI value from both methods. Moreover, bulk methods like zeta potential and IEF suffer not only from the suspended contamination within the bulk solution but also from the applied potential gradient-dependent charge mobility, which may induce distortion in the studied systems. This can be easily avoided with a single-molecule technique like CFM. Worth to mention that both of the mentioned methods can’t avoid the contribution from aggregation and hence looses their true measurements. Since the described method is single molecule-based and free from any external force which can distort the surface charge distribution of the studied system, we can measure the pI most accurately for any zwitterionic system (amino acids, peptides, proteins, cells, pathogens, etc.) including infectious viruses.

### Theoretical pI calculation from the outer region of E-protein

The theoretical Isoelectric point (pI) of the outer region for DENV-2 and DENV-4 envelope proteins obtained from the sequence level information from the ProtParam server^37^ were estimated to be 6.82 ± 0.25 and 6.75 ± 0.33, respectively (**Table S2**). The overall charge on the extracellular domain of the E protein ranges from mostly neutral to slightly negative (**Table S2**). Sequence-based pH-titration curves suggest similar pI values for the E proteins of DENV-2 (pI = 6.8 ± 0.18) and DENV-4 (pI = 6.75 ± 0.23) (**Figure 5 a, b, Table S2**). However, the sequence-based pI estimation of the E protein (pI ~ 6.8) differs from the much lower pI value estimated from the experiments. Clearly, the sequence-based pH-titration curve does not consider the folded structure, where a significant portion of the sequence remains buried inside the protein, and their charges are not accessible to the AFM probe.^32^ The protonation states of titrable amino acid side chains strongly depend on the local environments^38,39,40,41,42,43^, which is ignored in the sequence-based pH titration curves^44^ (**Figure 5 a, b**).

**Figure 5.**
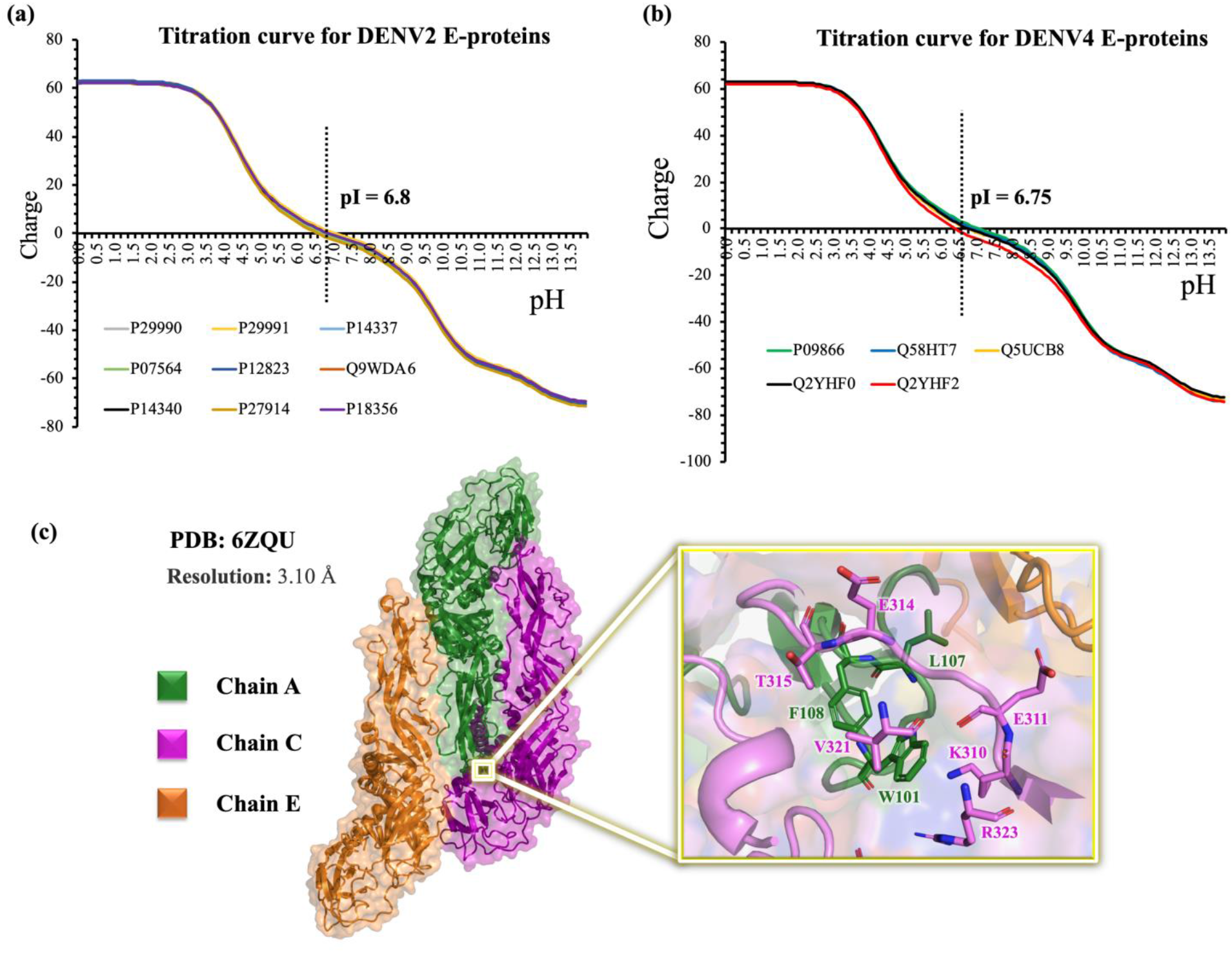
Theoretical titration curves of different strains of a) DENV-2 and b) DENV-4 envelope protein from its sequence information. (c) The structure of trimeric DENV-2 E-protein derived from PDB 6ZQU. The zoomed-in view of three important residues for trimerization (W101, L107, and F108) and their surrounding residues at the trimer interface have been shown in the yellow box.

To obtain insights into the protein’s surface charge distribution and the altered protonation states of titrable residues due to local environments, the Cryo-EM structure of DENV-2 surface protein (PDB 6ZQU^45^) was used as a template, and the 3D structure was subjected to pI estimation. The structure (PDB 6ZQU^45^) contains three chains of Envelope proteins (chains A, C, and E) that constituted the viral surface in a trimeric arrangement (**Figure 5c**). Three hydrophobic residues, W101, L107, and F108 (**Figure 5c**), were reported to be crucial for the trimerization of E protein at acidic pH to form the fusion loop^17,46^. These residues are surrounded by titrable residues like E311 and E314, which can alter their protonation states upon pH change (**Figure 5c, Figure 6**).

**Figure 6.**
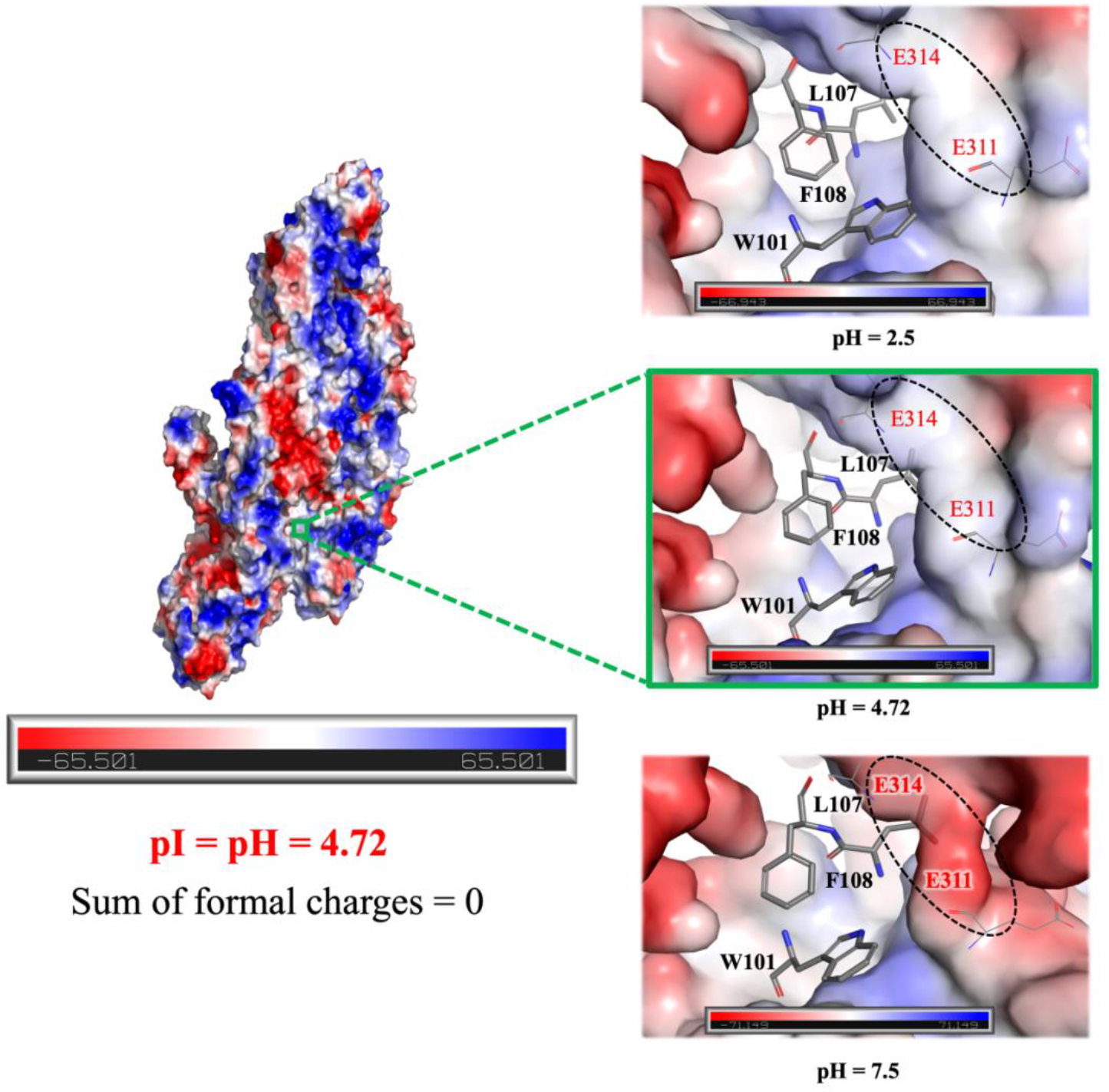
Electrostatic potential map of DENV-2 surface E-protein trimer at estimated theoretical pI (pH = 4.72). Zoomed-in view of the electrostatic potential map of regions surrounding trimerization promoting W101, L107, and F108 residues over different pHs have shown on the right side. Positive to negative charge is shown in blue to red scale while white regions denote regions of neutral charges. (Created in PyMOL).

The electrostatic potential maps of the DENV-2 envelope protein (**Figure S6**) provided the pattern of surface charge distribution on the viral envelope. The sum of the formal charge of individual structures at different pHs provided an estimate of how the overall charge on the protein changes upon changing the pH (protonation states). As expected, low pH increases the positive charge in the envelope protein (**Figure S6**). Whereas, an increase in the pH decreases the net positive charge of the envelope protein due to the deprotonation of several titrable amino acids (**Figure S6**). Interestingly, at pH = 4.72 (**Figure 6, Figure S6**), the protein’s net charge was estimated to be zero, and thus this pH serves as the isoelectric pH (pI) of the viral surface protein. The structure-based pI estimation was found to be closer to the experimentally determined pI relative to the conventional sequence-based pI estimation (**Figure 5 a, b**). Note, the 3D template structure did not include the chemical modifications (viz., glycosylation, acetylation, and phosphorylation, etc.)^32,47^ and was limited in modeling the real complex architecture of the viral envelope (including various proteins, lipids, cholesterol, and nucleic acids)^48^. These modifications will undoubtedly alter the overall charge on the viral envelope and justify the deviation between the experimental and structure-based pI estimation. Despite a slight deviation, it is clearly demonstrated that the structure-based titration curve estimation is superior to the conventional sequence-based approach.

The pKa of multiple titrable residues in the 3D structures of the viral surface E-protein (marked in “*”, **Figure S7**) was estimated to be distinctly different relative to their free amino acid form. For example, the side-chain pKa of E195 (chain E), E202 (chain C), and D22 (chain A) were estimated to be pKa > 7; thus, these residues were likely to be protonated (i.e., neutral) even at pH = 7. Note, with a pKa of 3.9 and 4.3, aspartic acid and glutamic acid, respectively^49^ have been classified as negatively charged amino acids in water. On the other hand, some aspartate and glutamate residues in the 3D structure have significantly low pKa, and hence they retain their negative charge even at very low pH. Interestingly, a drastic pKa shift towards lower value was evident (**Figure S7**) for some of the buried histidine residues (viz., H27, H244 (chain A, C), H261 (chain A, C), H282 (chain A, C, E), and H317 (chain A, C, E)). These buried histidines are likely to be neutral due to their hydrophobic environment. Very low pKa of K88 (Chain E) indicates that K88 will tend to lose its positive charge in pH as low as ~5. Clearly, the microenvironment around the titrable residues in the folded protein determines the pKa, thus, the overall charge on the protein. An increase in hydrophobicity and the proximity of charged residues are known to alter the protonation state, which is linked with the pKa shift in the folded protein structures^38,39,40^. In fact, more than 50% of the titrable residues were identified to be buried in the protein structure, which led to changes in pKa (**Figure S8**), and hence the overall charge of the protein in its folded form differs from the ones obtained only from sequence information. The E-proteins have been reported to trimerize in acidic pH, which involves three key residues W101, L107, and F108^50,7^. The negative charges of the environment around these three residues were found to decrease (from negatively charged to relatively neutral) at acidic pH (**Figure 6**). Protonation of the neighboring glutamic acid residues (E311 and E314) at the lower pH is primarily responsible for the same (**Figure 6**). The neutralization of the neighboring E311 and E314 residues at lower pH might stabilize the hydrophobic residues W101, L107, and F108 and facilitate trimerization. The overall negative change on E311 and E314 at high pH might disfavor the correct orientation and accommodation of the hydrophobic residues (W101, L107, and F108). Thus, it seems that the overall negative charge (in the pocket hosting W101, L107, and F108 residues) at higher pH might alter the local environment and is a possible reason for the destabilization of the trimeric complex.

In conclusion, we have reported the isoelectric point (pI) of two different infectious dengue serotypes, DENV-2 and DENV-4, respectively. Though the measured pI values for DENV-2 (4.05) and DENV-4 (3.94) are almost close to each other, thus it can be predicted from our study that the infectivity depends more on the structural orientation of E protein during pathogenesis over the cumulative charge of the virus. It is interesting to speculate that minor differences in virion surface charge distribution may lead to a significant difference in the pH at which the transition from dimer to trimer is induced in the E-protein. Such a temporal difference may have implications on the rate of virus entry into the host cell, with further effect on the kinetics of viral replication. Our single-particle CFM study and the associated theoretical analysis reveal the significance of the hydrophobic fusion loop and surrounded residues of E proteins which binds with the desired receptor in a pH-dependent manner followed by endosomal clearance. Other than surface proteins of DENV, it is known that the infection of all four serotypes is highly regulated by their non-structural (NS) proteins, like NS5 of DENV-4 has low nucleus localization efficiency than other serotypes. Also, the structural differentiation of NS1 among serotypes governs the endothelial barrier permeability^51,52^. Keeping all these previous studies in mind we can appreciate that the infectivity of DENV serotypes is crucially guided by their inclusive genetic changes which may not drastically change the cumulative surface charge of a whole virus but the domain-specific local charge distribution helping the different pockets of a protein to elicit their activity.

## Table of Content

The single-vesicle level infectivity of DENV serotypes is crucially guided by their inclusive genetic changes which may not drastically change the cumulative surface charge of a whole virus but the domain-specific local charge distribution helps the different pockets of a protein to elicit their activity.

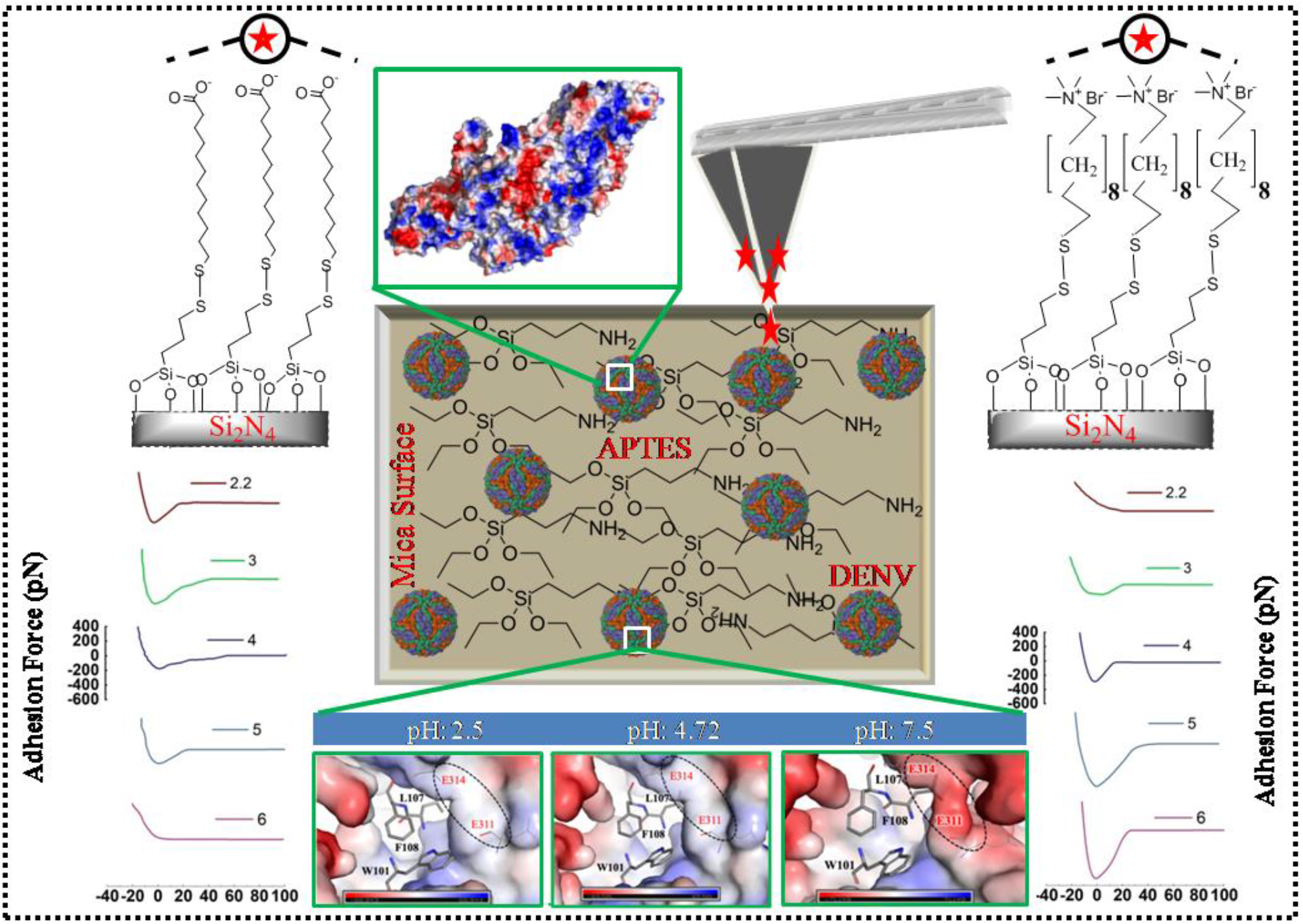

## Notes

### Competing Interest Statement

The authors have declared no competing interest.

