## Supplementary Information Document for "E-Protein Protonation Titration-induced Single Particle Chemical Force Spectroscopy for Microscopic Understanding and pI Estimation of Infectious DENV"

**EXPERIMENTAL SECTION:**

Chemicals: Sodium chloride (NaCl), citric acid monohydrate (HOC(CH_2_COOH)2(COOH).H_2_O), sodium citrate tribasic dihydrate (HOC(COONa)(CH_2_COONa)_2_), 12-mercaptododecanoic acid (HS(CH_2_)_11_COOH), APTES (Si(OCH_2_CH_3_)_3_(H_2_NCH_2_CH_2_CH_2_)), MPTMS (HS CH_2_CH_2_CH_2_Si(OCH_3_)_3_), DTT(HSCH_2_CH(OH)CH(OH)CH_2_SH), Isopropanol (CH_3_CH(OH)CH_3_), 11-mercaptoundecyl-N,N,N-trimethylammonium bromide (HS(CH_2_)_11_N(CH_3_)_3_Br), sodium hydroxide (NaOH), sodium phosphate monobasic (NaH_2_PO_4_) and sodium phosphate dibasic (Na_2_HPO_4_) were purchased from Sigma-Aldrich. All aqueous solutions or buffers were prepared using purified water with a resistivity of ≥15 MΩ·cm from a Milli-Q filtration system and filtered with a 0.2 µm syringe filter before use. Citrate buffer (CB) solutions with different pH (3.0–6.0) were prepared by mixing different volume percentages of 20 mM stock solution of citric acid and sodium citrate tribasic. Phosphate buffer (PB) solution at pH 7.4 was prepared by mixing 20 mM stock solution of sodium phosphate monobasic and sodium phosphate dibasic. The final pH was adjusted with 1M NaOH or HCl, as needed, and measured with a calibrated benchtop pH meter (Oakton, pH 700).

**Preparation of buffers:**

20 mM PBS was prepared at pH 7.4 and 20 mM phosphate-citrate buffer was prepared at pH 2.2 by mixing 0.01 g disodium hydrogen phosphate (MW: 141.96 g/mol) and 0.41 g citric acid (MW: 210.14 g/mol) in 20 mL water. Other 20 mM citrate buffer (CB) solutions were prepared from stock solutions of  0.02 M citric acid and 0.02 M trisodium citrate (TSC) at pH 2.2.0 - 6.0 as shown in **table S1**^1^.

**Table S1: Preparation of citrate buffer at pH 3 to 6:**

| pH of CB | Volume of TSC (0.02M) | Volume of Citric acid (0.02M) |
| --- | --- | --- |
| 3 | 9.0 | 41.0 |
| 4 | 20.5 | 29.5 |
| 4.5 | 26.5 | 23.4 |
| 5 | 32.5 | 17.5 |
| 5.5 | 38.3 | 11.6 |
| 6 | 44.2 | 5.7 |

All buffers were prepared in autoclaved Milli-Q® water and passed through 25 mm Whatman filter paper before adjusting the final pH with a calibrated pH meter with concentrated HCl and NaOH.

**Preparation of Inactivated virus samples:**

The inactivated dengue virus (DENV) samples were prepared as mentioned before by De et al.^2^ In brief, 4-aminomethyltrioxsalen hydrochloride is added to 10 mL of viral supernatant to achieve the final concentration of 10 mg/mL. The supernatant was then irradiated with a UV torch (365 nm, UVP UVLMS-38EL series 3UV lamp Upland CA, USA, 8-Watt 230V-50Hz, 0.16 Amps) after its distribution by 2 mL in 6 well polystyrene plates of diameter 3.5 cm (Corning™ Costar™) for 20-25 minutes.

**AFM probe functionalization with a charged chemical group:**

At first, the surface of the AFM probe (SNL-10) was washed with acidic water and modified with an organo-silane monolayer by emerging it into a 10 mM of 3-mercaptopropyl trimethoxysilane (3-MPTMS) in isopropanol (in presence of 5 mM DTT) for a period of 12 h. After that, the probes were taken out from the 3-MPTMS solution for further modification followed by bath-sonication in acetone and ethanol for 2-3 minutes.^3^

**Positively and negatively charged AFM probe modification:**

The charge modification of the 3-MPTMS monolayer protected probes was carried out by the following reported protocol.^1^ For the preparation of negatively-charged probes, they were first emerged in a 4 mM solution of 12-mercaptododecanoic acid in ethanol for 24 h, washed then with ethanol, and dried in a desiccator. On contrary, positively charged probes were prepared by immersing probes in a 10 mM solution of 11-mercaptoundecyl-N,N,N- trimethylammonium bromide in ethanol for 48 h, washed then with ethanol and dried in a desiccator. On each consecutive experiment, we made a freshly prepared functionalized probe. A detailed functionalization procedure and its chemical reactions are shown in **Figure S1** where as the magnified SEM images of these modified probes are shown in **Figure S2**.

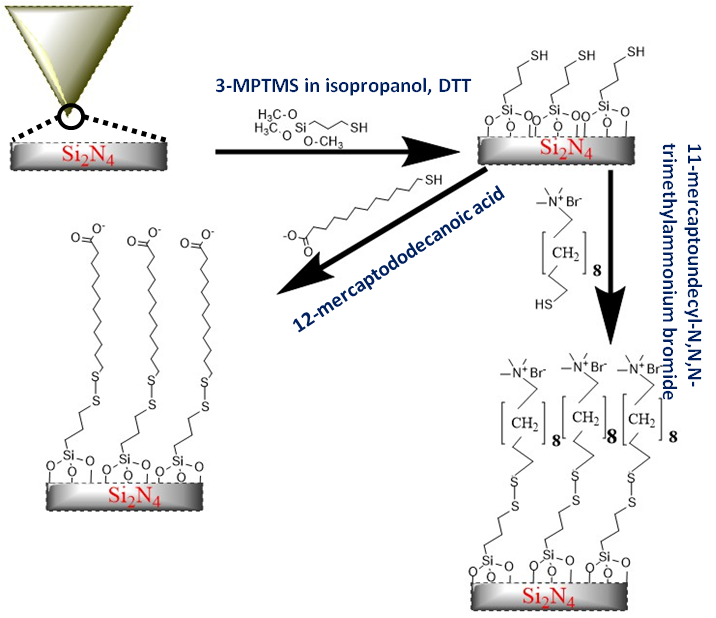

**Figure S1. Modification of AFM probe:** At the first step the SNL10 probe is modified with 10 mM (3-mercaptopropyl) trimethoxysilane (3-MPTMS) .In the second step, the probe is modified either with 12-mercaptododecanoic acid or 11-mercaptoundecyl-N,N,N-trimethylammonium bromide for negative and positive charge modification respectively.

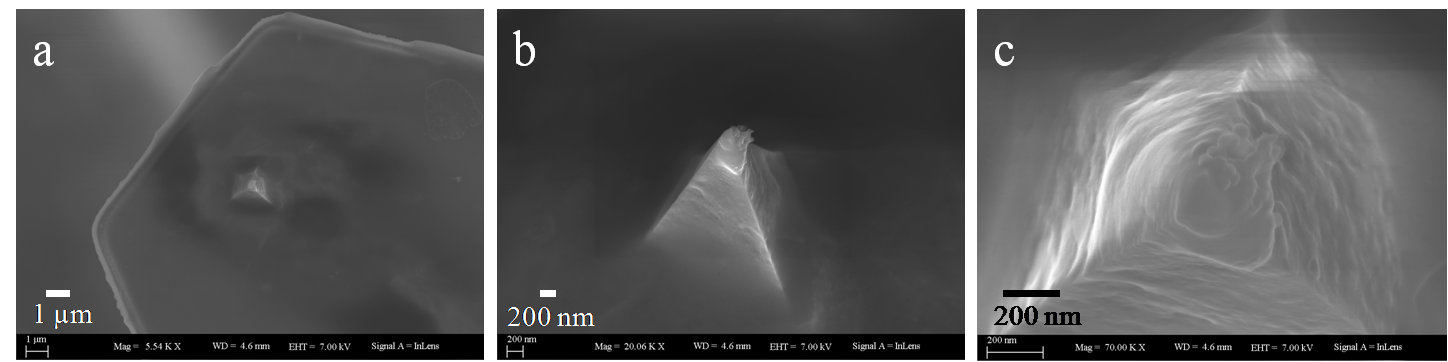

**Figure S2. SEM of modified probe:** The (a) tip, (b) side, and (c) top view of Bruker’s SNL10 probes after 2-step modification with MPTMS and mercaptododecanoic acid to get a negatively charged tip.

**Preparation of AFM Sample:**

Freshly cleaved muscovite mica (SPI Supplies, West Chester, USA) was installed on a fresh glass slide using sticky glue. 50 µL of 10^4^ PFU/mL DENV samples (psoralen inactivated) were incubated on mica and kept at room temperature for 20 minutes. It was then washed gently two times with 100 µL of 20 µM CB or PBS of the desired pH and the whole setup was placed on the AFM stage. Again, the mica was rehydrated before the experiment with 100 µL CB or PBS of the desired pH. For virus film preparation, the mica surface was first incubated with 0.0001% (3-aminopropyl) triethoxysilane (3-APTES) for 30 min and rinsed 3 times in 100 µL of Milli-Q water. 70 µL of 10^4^ PFU/mL DENV solutions were then drop casted on modified mica and placed at 4°C overnight. The next day, the mica was taken out and rinsed with buffer, and placed on the AFM stage after rehydration with the desired buffer for liquid imaging and force spectroscopy experiments.

**AFM imaging and analysis:**

All AFM experiments were performed in liquid with tapping and contact mode at room temperature on a BrukerBioScope Catalyst Atomic Force Microscope instrument. AFM topographic images were obtained using tapping mode in CB or PBS with a Bruker SNL-10 cantilever with a spring constant of k = 0.03 N/m and a frequency range of 12-18 kHz. The tip was engaged at a minimum scan range condition in the feedback loop condition to prevent tip contamination. The data analysis of surface topography images was performed using the Nasoscope analysis software.

**Chemical force measurement and analysis:**

All AFM force measurements were performed in contact mode. The spring constant of both positively and negatively charged modified AFM probes was calibrated before the force measurement by using the thermal noise method. All the adhesion force experiments are measured on the DENV film sample and with reference to the retract curve of the ramp and the Force-volume (F-V) method. For each set of experiments, at least 500  F-D curves and a minimum of 100 ramp curves are taken and each experiment was repeated three times with a separate set of the modified probe and DENV film sample. Individual adhesion histograms and representative force-distance curves for DENV-2 and DENV-4 with negatively and positively charge terminated probes under varying pH at 2.2, 3, 4, 5, and 6 are shown in **Figures S3** and **S4**.

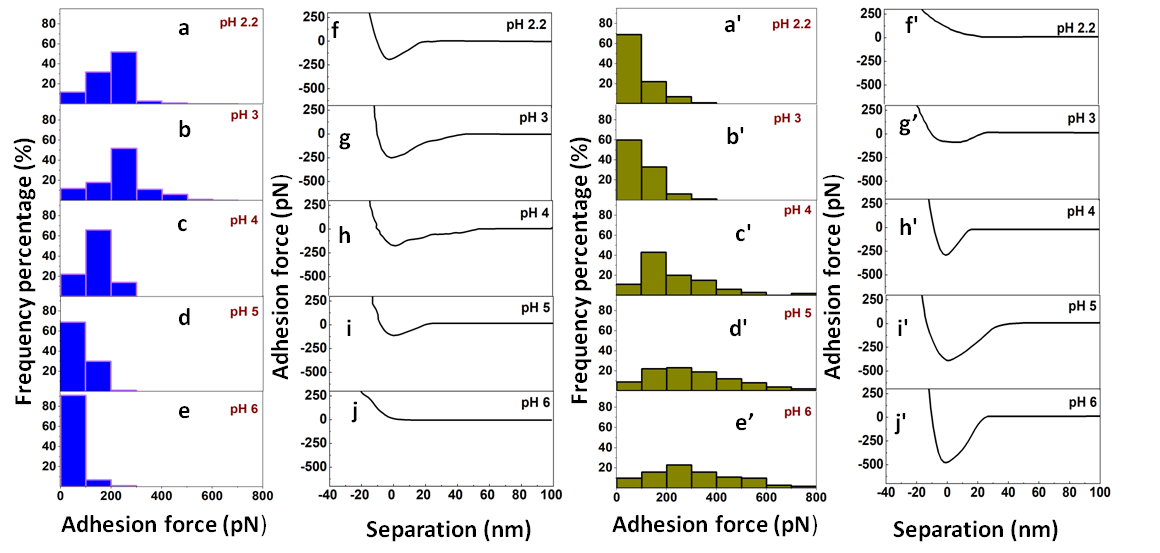

**Figure S3: Individual adhesion histograms and representative force-distance curves**: Adhesion forces of DENV-2 with negatively (a-e) and positively (a′-e′) charge terminated probe under varying pH at 2.2, 3, 4, 5, and 6. Corresponding retract curves for negatively (f-j) and positively (f′-j′) charged probe.

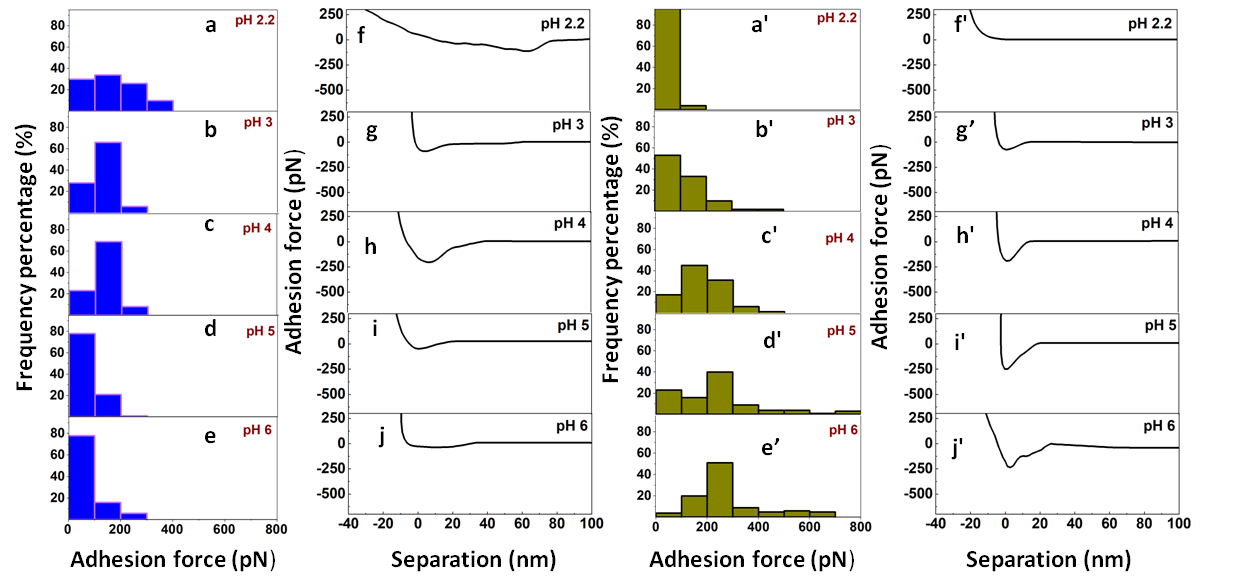

**Figure S4. Individual adhesion histograms and representative force-distance curves**: Adhesion forces of DENV-4 with negatively (a-e) and positively (a′-e′) charge terminated probe under varying pH at 2.2, 3, 4, 5, and 6. Corresponding retract curves for negatively (f-j) and positively (f′-j′) charged probe.

Force measurements and curves were taken in a 20 mM CB at different pHs. Before each force measurement, the sample was imaged under a buffer with an unmodified SNL10 tip to optimize the region of interest and F-V curves were taken with the functionalized cantilever in the same area. Five sets of F-V curves were taken for each modified tip and adhesion forces were analyzed with the BrukerNanoscope Analysis software. To find out the pI of the DENV, the mean adhesion force from all curves was plotted as a function of pH and further fit a sigmoidal curve as described by Miet al.^1,4^

$$Y=\frac{y_{f}}{1+e^{-\frac{\left( x-x_{0} \right)}{\tau}}}$$

where Y is the adhesion force, x is the set pH, x_0_ is the measured pI, y_f_ is the maximum adhesion force, and the rate constant for the change of the mean adhesion force as a function of pH is given by 1/τ.

**Zeta potential measurement:**

Inactivated DENV with a concentration of 10^4^ PFU/mL was diluted to 1:500 in a 20 mM CB at the desired pH. The zeta potential was measured using a Malvernmade Zetasizer ZS90 instrument at 25°C using a capillary zeta cell (750 μL) with an equilibration time of 100s.

**Computational Methods:**

**Sequence Retrieval and Identification of conserved regions:**

Sequence information for different strains of dengue virus serotype 2 (DENV-2; accession number: P29990, P29991, P14337, P07564, P12823, Q9WDA6, P14340, P27914, and P18356) and serotype 4 (DENV-4; accession number: P09866, Q58HT7, Q5UCB8, Q2YHF0, and Q2YHF2) were retrieved from the UniProt Knowledgebase (UniProtKB) database^5^. The amino acid sequences corresponding to the Envelope (E) protein of the viruses (**Table S2**) were extracted using the ExtractAlign tool of EMBOSS explorer^6^ (https://www.bioinformatics.nl/cgi-bin/emboss/extractalign). The different domains present in the E protein (of DENV-2 and DENV-4) were identified from the InterPro^7^ and Conserved domain database (CDD) server^8^. The extent of the evolutionary conservation of the E-protein residues was accessed from Multiple Sequence Alignment (MSA) implemented inClustal Omega server^9^.

**Table S2.** Physiochemical characterization and domains in different strains of DENV2 and DENV4 E-proteins.

| **DENV-2 strains** | | | | | | | | | | |
| --- | --- | --- | --- | --- | --- | --- | --- | --- | --- | --- |
| **UniProt ID** | **E-protein position** | **Length** | **Molecular weight (Da)** | **Theoritical pI** | | **Grand average of hydropathicity (GRAVY)** | **Total number of negatively charged residues (Asp + Glu)** | **Total number of positively charged residues (Arg + Lys)** | **Domains** | |
|  |  |  |  | **ProtParam** | **Prot Pi** |  |  |  | **positions in E protein** | **Domain name** |
| P29990 | 281-725 | 445 | 49145.57 | 7.16 | 7.047 | -0.287 | 50 | 50 | 2-296 | Flavivirus glycoprotein, central and dimerisation domains |
|  |  |  |  |  |  |  |  |  | 298-394 | Flavivirus glycoprotein, immunoglobulin-like domain |
|  |  |  |  |  |  |  |  |  | 399-445 | Flavivirus envelope glycoprotein E, stem/anchor domain |
| P29991 | 281-725 | 445 | 49153.62 | 7.16 | 7.047 | -0.269 | 50 | 50 | 2-296 | Flavivirus glycoprotein, central and dimerisation domains |
|  |  |  |  |  |  |  |  |  | 298-394 | Flavivirus glycoprotein, immunoglobulin-like domain |
|  |  |  |  |  |  |  |  |  | 399-445 | Flavivirus envelope glycoprotein E, stem/anchor domain |
| P14337 | 281-725 | 445 | 49070.48 | 6.86 | 6.85 | -0.261 | 50 | 49 | 2-296 | Flavivirus glycoprotein, central and dimerisation domains |
|  |  |  |  |  |  |  |  |  | 298-394 | Flavivirus glycoprotein, immunoglobulin-like domain |
|  |  |  |  |  |  |  |  |  | 399-445 | Flavivirus envelope glycoprotein E, stem/anchor domain |
| P07564 | 281-725 | 445 | 49098.58 | 6.65 | 6.678 | -0.244 | 51 | 49 | 2-296 | Flavivirus glycoprotein, central and dimerisation domains |
|  |  |  |  |  |  |  |  |  | 298-394 | Flavivirus glycoprotein, immunoglobulin-like domain |
|  |  |  |  |  |  |  |  |  | 399-445 | Flavivirus envelope glycoprotein E, stem/anchor domain |
| P12823 | 281-725 | 445 | 49047.4 | 6.86 | 6.849 | -0.285 | 51 | 50 | 2-296 | Flavivirus glycoprotein, central and dimerisation domains |
|  |  |  |  |  |  |  |  |  | 298-394 | Flavivirus glycoprotein, immunoglobulin-like domain |
|  |  |  |  |  |  |  |  |  | 399-445 | Flavivirus envelope glycoprotein E, stem/anchor domain |
| Q9WDA6 | 281-725 | 445 | 49058.43 | 6.48 | 6.528 | -0.266 | 52 | 49 | 2-296 | Flavivirus glycoprotein, central and dimerisation domains |
|  |  |  |  |  |  |  |  |  | 298-394 | Flavivirus glycoprotein, immunoglobulin-like domain |
|  |  |  |  |  |  |  |  |  | 399-445 | Flavivirus envelope glycoprotein E, stem/anchor domain |
| P14340 | 281-725 | 445 | 49036.47 | 6.86 | 6.85 | -0.257 | 50 | 49 | 2-296 | Flavivirus glycoprotein, central and dimerisation domains |
|  |  |  |  |  |  |  |  |  | 298-394 | Flavivirus glycoprotein, immunoglobulin-like domain |
|  |  |  |  |  |  |  |  |  | 399-445 | Flavivirus envelope glycoprotein E, stem/anchor domain |
| P27914 | 1-445 | 445 | 49086.48 | 6.48 | 6.52 | -0.26 | 52 | 49 | 2-296 | Flavivirus glycoprotein, central and dimerisation domains |
|  |  |  |  |  |  |  |  |  | 298-394 | Flavivirus glycoprotein, immunoglobulin-like domain |
|  |  |  |  |  |  |  |  |  | 399-445 | Flavivirus envelope glycoprotein E, stem/anchor domain |
| P18356 | 181-625 | 445 | 49056.46 | 6.86 | 6.85 | -0.262 | 50 | 49 | 2-296 | Flavivirus glycoprotein, central and dimerisation domains |
|  |  |  |  |  |  |  |  |  | 298-394 | Flavivirus glycoprotein, immunoglobulin-like domain |
|  |  |  |  |  |  |  |  |  | 399-445 | Flavivirus envelope glycoprotein E, stem/anchor domain |
| Average | - | 445 | 49083.72 | 6.82 | 6.803 | -0.2656 | 50.66 | 49.33 | - | - |
| **DENV-4 strains** | | | | | | | | | | |
| P09866 | 280-725 | 446 | 48709.7 | 7.18 | 7.064 | -0.213 | 49 | 49 | 2-296 | Flavivirus glycoprotein, central and dimerisation domains |
|  |  |  |  |  |  |  |  |  | 298-394 | Flavivirus glycoprotein, immunoglobulin-like domain |
|  |  |  |  |  |  |  |  |  | 399-445 | Flavivirus envelope glycoprotein E, stem/anchor domain |
| Q58HT7 | 280-723 | 444 | 48527.59 | 6.89 | 6.874 | -0.188 | 50 | 49 | 2-296 | Flavivirus glycoprotein, central and dimerisation domains |
|  |  |  |  |  |  |  |  |  | 298-394 | Flavivirus glycoprotein, immunoglobulin-like domain |
|  |  |  |  |  |  |  |  |  | 399-445 | Flavivirus envelope glycoprotein E, stem/anchor domain |
| Q5UCB8 | 280-723 | 444 | 48464.32 | 6.68 | 6.707 | -0.216 | 50 | 48 | 2-296 | Flavivirus glycoprotein, central and dimerisation domains |
|  |  |  |  |  |  |  |  |  | 298-394 | Flavivirus glycoprotein, immunoglobulin-like domain |
|  |  |  |  |  |  |  |  |  | 399-445 | Flavivirus envelope glycoprotein E, stem/anchor domain |
| Q2YHF0 | 280-725 | 446 | 48727.72 | 6.71 | 6.736 | -0.205 | 50 | 48 | 2-296 | Flavivirus glycoprotein, central and dimerisation domains |
|  |  |  |  |  |  |  |  |  | 298-394 | Flavivirus glycoprotein, immunoglobulin-like domain |
|  |  |  |  |  |  |  |  |  | 399-445 | Flavivirus envelope glycoprotein E, stem/anchor domain |
| Q2YHF2 | 280-725 | 446 | 48790.58 | 6.29 | 6.355 | -0.224 | 52 | 47 | 2-296 | Flavivirus glycoprotein, central and dimerisation domains |
|  |  |  |  |  |  |  |  |  | 298-394 | Flavivirus glycoprotein, immunoglobulin-like domain |
|  |  |  |  |  |  |  |  |  | 399-445 | Flavivirus envelope glycoprotein E, stem/anchor domain |
| Average | - | 445.2 | 48643.98 | 6.75 | 6.75 | -0.2092 | 50.2 | 48.2 | - | - |

**Theoretical pI calculation of DENV-2 and DENV-4 E proteins:**

Expasy’s ProtParam server^10^ was used to estimate the theoretical pI of the envelope proteins based on its sequence information. The sequence information-based pH versus charge titration curves was determined by the ProtPIProtein tool (www.protpi.ch/Calculator/ProteinTool). To understand the 3D surface charge distribution of the DENV-2 envelope protein, a template Cryo-EM structure of the mature Dengue virus-2 (PDB 6ZQU^11^, resolution= 3.1 Å) was obtained from the Protein Data Bank (PDB)^12,13^ ([www.wwpdb.org](http://www.wwpdb.org)) and the protonation state of the titratable amino acid residues in the 3D structure (PDB 6ZQU)^11^ at various pH was estimated by the PROPKA^14^ analyzer implemented in Schrodinger Maestro (Academic version 2022-4; Schrödinger). The sum of the formal charges of the 3D structure at various pH was estimated in PyMOL^15^. The surface electrostatic potential map on the 3D structure of the E-protein in a vacuum was estimated and visualized using PyMOL^15^ software.

**Sequence alignment, conserved domain, and physicochemical properties:**

From the mentioned UniPort database, the outer region of the E protein of the DENV-2 and DENV-4 strains are 445 (molecular weight of ~49 kDa) and 444 - 446 amino acid residues long (molecular weight ~48 kDa) respectively. Among the three conserved functional domains, E-I and E-II belong to a class-II fusion protein which facilitates host-receptor binding and fusion. These domains (I and II) in low pH (< 6.3) have been reported to undergo conformational changes and trimerization^13,14^. The trimeric form of E protein was reported to enhance the virus’s infectivity^13,15^. The E-III corresponds to a C-terminal stem region connected to two transmembrane helices. The negative value of the grand average of hydropathicity (GRAVY, **Table S2**) indicated the hydrophilic nature of the E protein. However, the E protein of DENV-2 is relatively more hydrophilic than DENV-4 (**Table S2**). High sequence conservation of the E proteins among the retrieved sequences was evident from multiple sequence alignment (**Figure S5**). Particularly, high conservation of the charged amino acid residues across different strains of DENV-2 and DENV-4 indicated that the E-proteins of different strains contain similar charge distribution for the sequence-specific aspect.

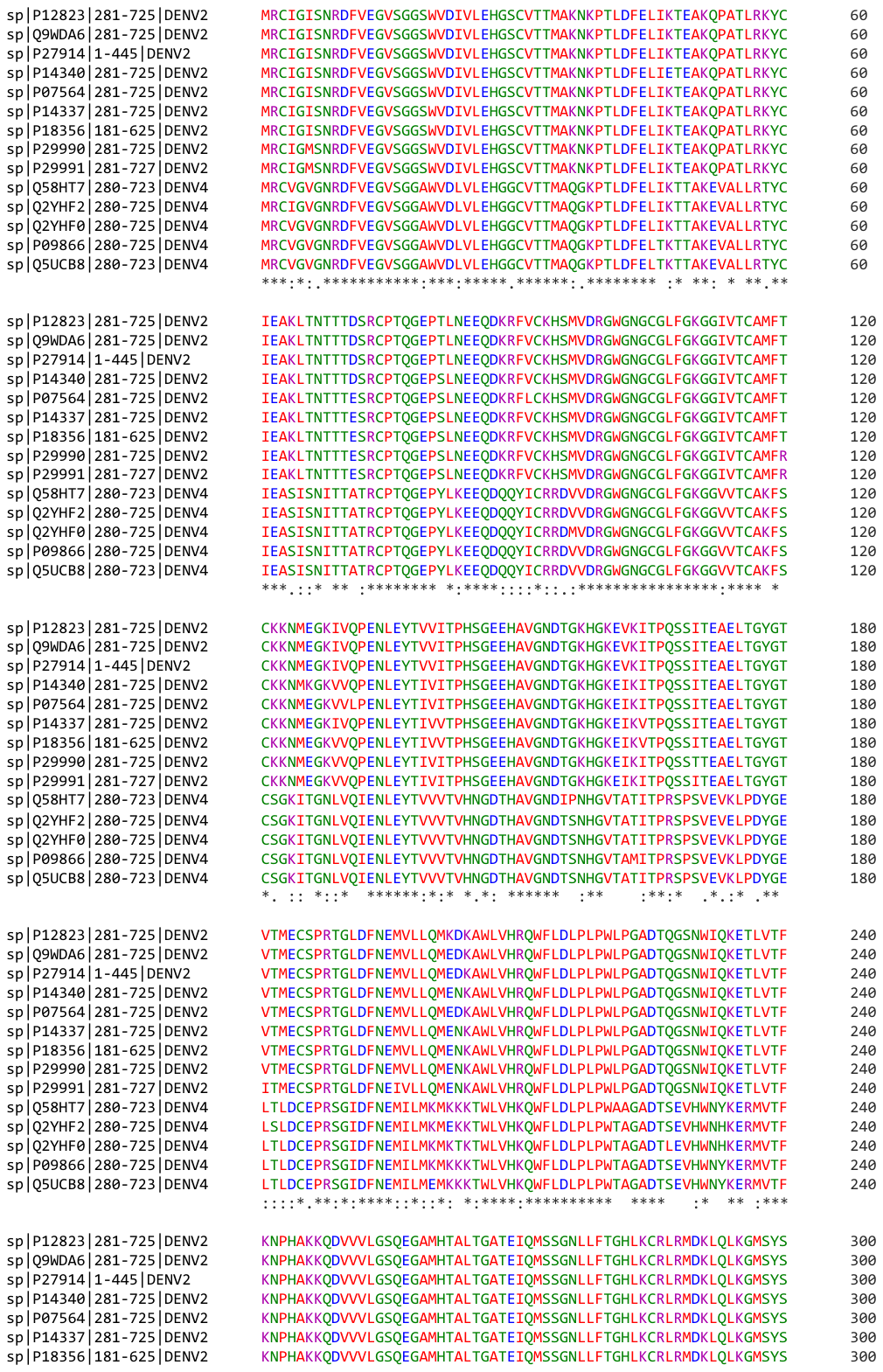

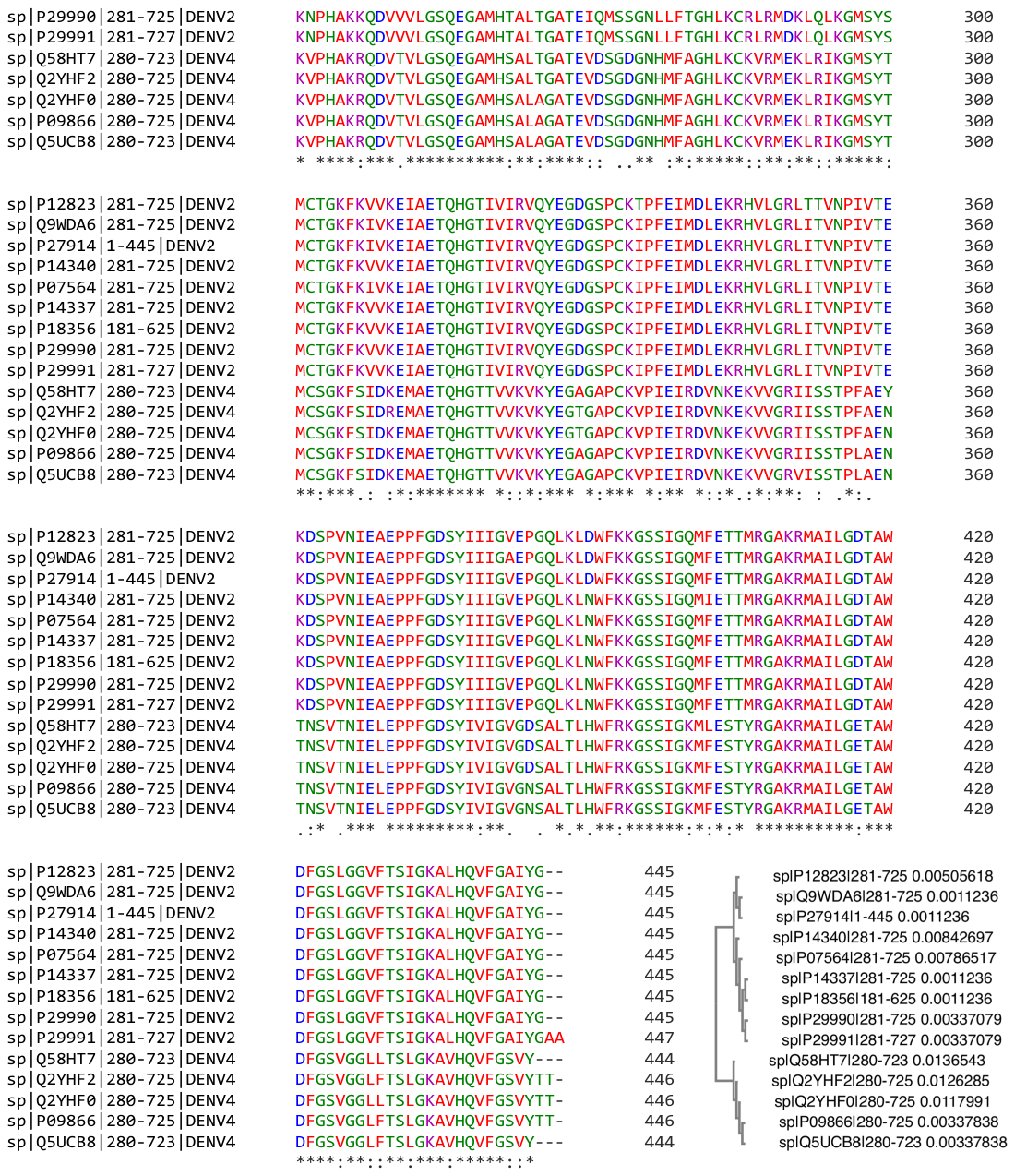

**Figure S5**. Multiple sequence alignment envelope (E) protein extracellular domains from different DENV-2 and DENV-4 strains. The guide tree demonstrates the DENV-2 and DENV-4 clusters. Asterisks (*) indicate fully conserved residues, colons (:) denote amino acid substitution with similar physiochemical properties, and dots (.) represent the substitution of amino acids with different physiochemical properties.

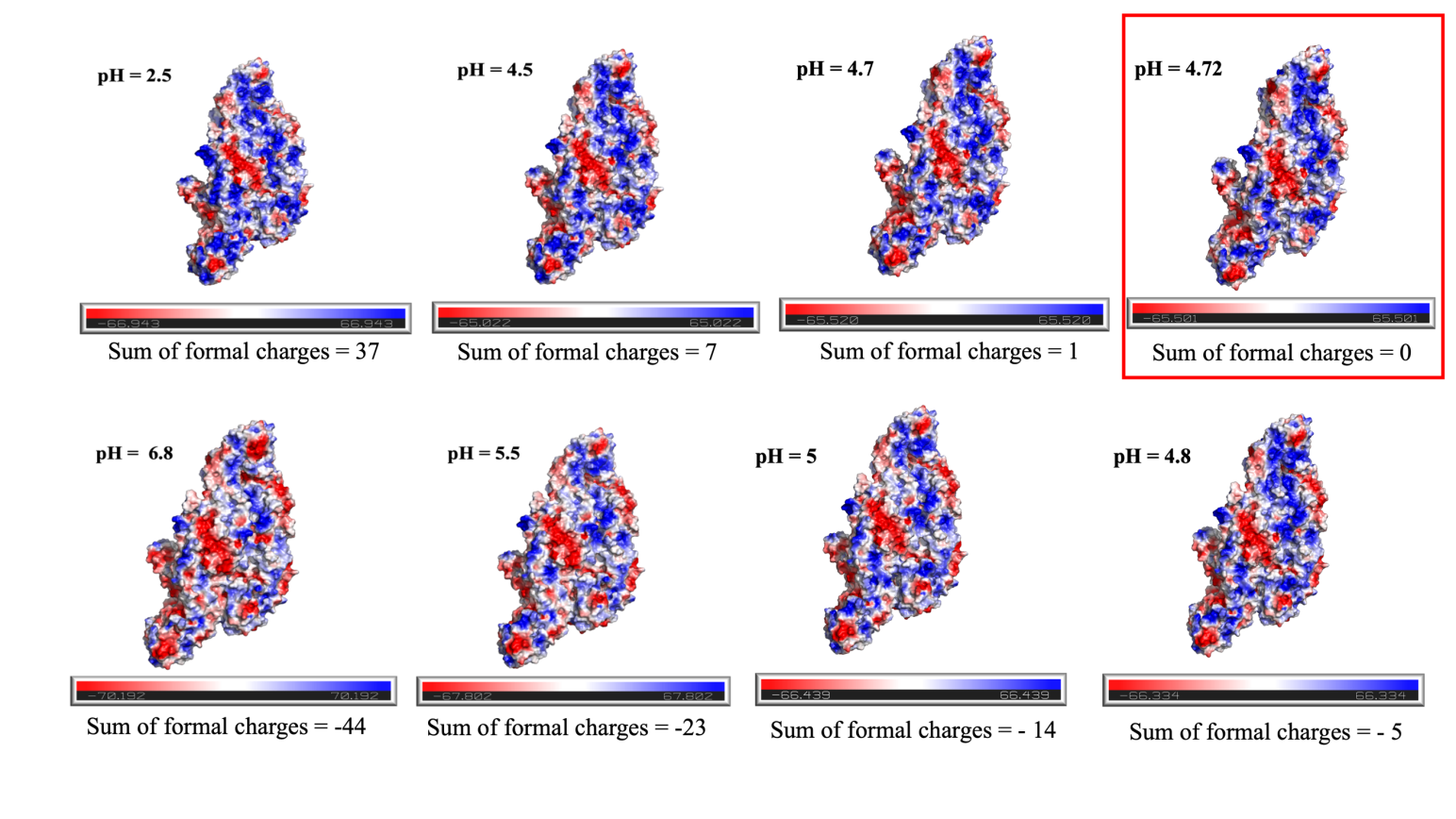

**Figure S6**. The electrostatic potential map of DENV-2 envelope protein at different pH (pH = 2.5, pH = 4.5, pH = 4.7, pH = 4.72, pH = 5, pH = 5.5, and pH = 6.8). The sum of the formal charges became zero at pH = 4.72 (estimated pI), marked in a red box. Positive to negative charge is shown in blue to red scale. White regions denote regions of neutral charges. (Created in PyMOL).

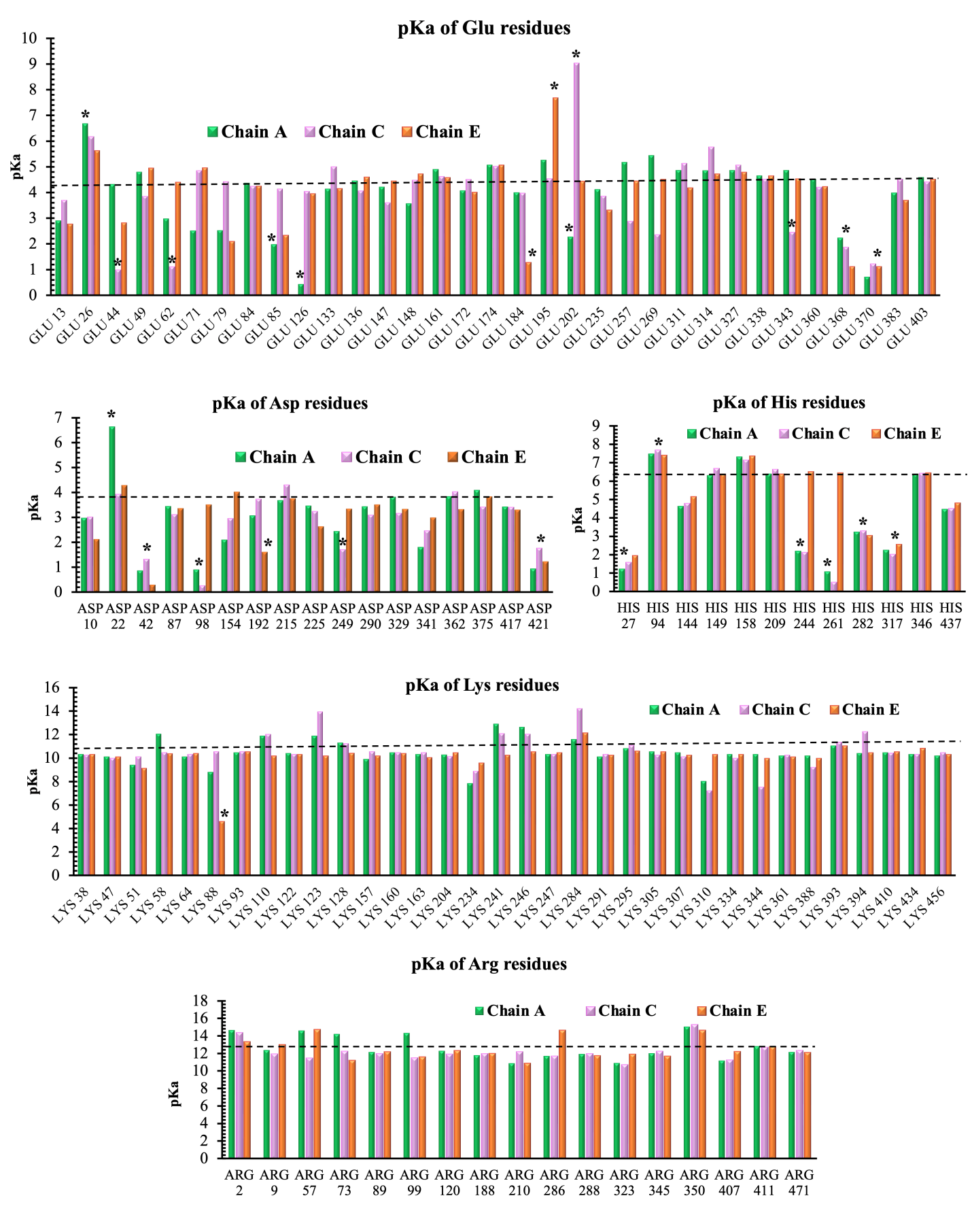

**Figure S7**. Estimated pKa for individual titrable residues in the DENV-2 surface E-protein structure (PDB: 6ZQU, chain A, C, and E) derived from PROPKA. The dashed line denotes the theoretical pKa of the amino acid side chains free in the water. Asterisks (*) denote residues with pKa highly deviated from their corresponding theoretical pKa.

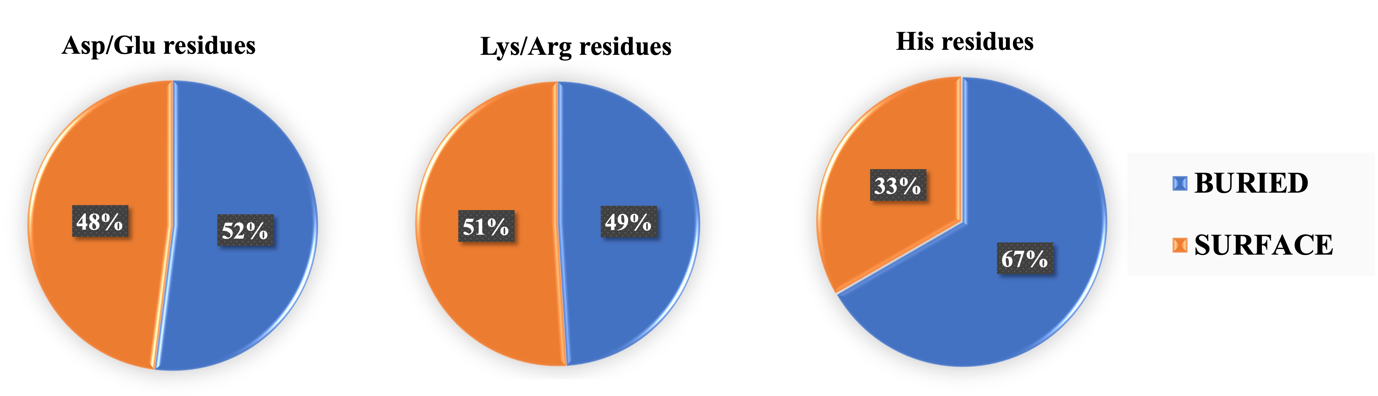

**Figure S8.** Percentage of titrable residues buried or present in the surface of DENV-2 E protein.
